# MolluscaGenes: A Transcriptomic Database for the Mollusca

**DOI:** 10.64898/2026.05.05.723003

**Authors:** Jorge L. Pérez-Moreno, Paul S. Katz

## Abstract

The phylum Mollusca constitutes one of the most taxonomically and morphologically diverse animal clades; however, the genomic exploration of this group has been hampered by fragmented and taxonomically incomplete transcriptomic resources. To address this fundamental limitation, we present MolluscaGenes, a centralized database that unifies transcriptomes from 299 molluscan species spanning all eight recognized classes, encompassing a broad array of tissues and developmental stages. MolluscaGenes provides searchable databases via BLAST and DIAMOND alongside a suite of 196 molluscan-optimized Hidden Markov Models (HMMs) for sensitive protein family identification. To demonstrate the utility of this resource, we performed a comprehensive phylum-wide characterization of the nicotinic acetylcholine receptor (nAChR) superfamily, recovering 3,586 sequences from over 190 species and resolving 15 distinct phylogenetic clades. This analysis revealed substantial lineage-specific expansions across multiple molluscan classes, the identification of novel clades with substitutions in canonical ligand-binding residues, and the evolutionary placement of chemotactile receptors (CRs) and CR-like sequences as predominantly cephalopod clades within the broader nAChR phylogeny. MolluscaGenes constitutes a foundational resource that will accelerate the elucidation of the unique biology and evolutionary history of Mollusca.

## 1. Introduction

Mollusca, the second-largest animal phylum after Arthropoda, encompasses over 70,000 described extant species distributed across eight recognized classes (Rosenberg 2014; Bouchet et al. 2016; Sigwart and Lindberg 2015). The evolutionary history of this phylum, spanning more than 500 million years since the Cambrian explosion, has produced an extraordinary breadth of body plans across marine, freshwater, and terrestrial ecosystems (Ponder et al. 2020). This taxonomic and morphological diversification renders Mollusca a fundamental system for investigating the genomic underpinnings of biomineralization, the independent elaboration of complex nervous systems, and the molecular basis of ecological adaptation across vastly different environmental regimes.

Despite this profound evolutionary significance, genomic research in molluscs has lagged behind other major animal phyla. Longstanding technical challenges, including the abundance of mucopolysaccharides that interfere with nucleic acid extraction and the prevalence of large, highly heterozygous, repeat-rich genomes, have historically complicated assembly efforts (Sun et al. 2020; Sigwart et al. 2021). Consequently, publicly available transcriptomic and genomic data remain fragmented across numerous repositories. The most comprehensive existing resource, MolluscDB 2.0, encompasses 88 high-quality genome assemblies (52 at chromosome-level) and transcriptomic datasets spanning all eight classes (Liu et al. 2021, 2025); however, deep transcriptomic coverage remains limited to approximately 60 species, predominantly bivalves and gastropods of economic importance. This taxonomic sampling is sparse relative to the estimated 70,000+ described species, with genome sequencing having historically favored economically valuable taxa or those with smaller, less repetitive genomes, leaving many evolutionarily significant lineages genomically unexplored (Sigwart et al. 2021).

The fragmentation of existing resources is compounded by a fundamental computational challenge: the detection of divergent protein homologs. Standard sequence similarity searches employing BLAST-based algorithms perform poorly when sequence identity falls below 30%, a common scenario when comparing deeply diverged molluscan lineages or when searching for homologs of vertebrate or insect proteins in lophotrochozoans (Terrapon et al. 2012). Profile Hidden Markov Models (HMMs), which incorporate position-specific information from multiple sequence alignments, offer substantially greater sensitivity for detecting distant homologs (Finn et al. 2016; Eddy 2011). Nevertheless, the HMM profiles available in databases such as Pfam are predominantly trained on vertebrate and ecdysozoan sequences, and may fail to detect divergent molluscan family members due to lineage-specific substitution patterns. This limitation proves particularly acute for rapidly evolving gene families, including sensory receptors and immune-related proteins (Tassia et al. 2021).

To address these challenges, MolluscaGenes was developed as a centralized database that unifies hundreds of newly assembled and previously published transcriptomes into a single, taxonomically comprehensive resource. A key methodological innovation is the multi-assembler transcriptome processing pipeline: studies have demonstrated that combining outputs from multiple de novo assemblers followed by intelligent redundancy reduction yields more complete assemblies than any single assembler alone (Nakasugi et al. 2014; Hsieh et al. 2019; Pérez-Moreno et al. 2023). The pipeline employs the nf-core/denovotranscript workflow, integrating Trinity and rnaSPAdes assemblies with EvidentialGene-based redundancy reduction to maximize coding sequence recovery while minimizing artifacts (Bhojwani et al. 2024; Gilbert 2019). Beyond assembly, MolluscaGenes incorporates a suite of molluscan-optimized HMMs developed through the TIAMMAt workflow (Tassia et al. 2021), which iteratively recalibrates generic Pfam profiles against the sequence diversity captured in the database. The resulting resource currently houses data from 299 species spanning all eight molluscan classes, with search capabilities including BLAST, DIAMOND (Buchfink et al. 2015), and optimized HMM-based protein family identification.

To demonstrate these capabilities, a phylum-wide characterization of the nicotinic acetylcholine receptor (nAChR) superfamily is presented as a case study. The nAChR superfamily, comprising members of the pentameric Cys-loop receptor family, mediates rapid synaptic transmission throughout the animal kingdom (Changeux 2012; Jones and Sattelle 2007). In molluscs, nAChRs have undergone remarkable evolutionary expansion, with bivalves harboring 99 to 217 paralogs compared with only 10 to 29 in vertebrates, accompanied by functional diversification into both cation- and anion-selective subtypes (Li et al. 2019; Dent 2006). Additionally, cephalopod chemotactile receptors (CRs) represent a derived clade within this superfamily, in which ancestral nAChRs have been neofunctionalized into contact-dependent chemosensors that enable octopuses to detect environmental stimuli through direct surface contact (Giesen et al. 2020; Allard et al. 2023). The breadth of taxonomic coverage in MolluscaGenes enables systematic investigation of the nAChR superfamily across the phylum, tracing the evolutionary trajectories from synaptic neurotransmission to environmental chemosensation and elucidating the genomic mechanisms underlying receptor diversification within and across molluscan classes.

## 2. Materials and Methods

### 2.1 Data Acquisition and Curation

Raw sequence data were gathered from publicly available sources, primarily the NCBI Sequence Read Archive (SRA). Searches were conducted in September 2024 using the query terms “Mollusca\[Organism\]ANDRNA-Seq\[Strategy\]OR Illumina\[Platform\]ORPacBio\[Platform\]” filtered for transcriptomic data. This search yielded candidate datasets, which were manually curated to remove metagenomic, environmental, and non-transcriptomic samples.

The selection strategy prioritized: (1) paired-end Illumina datasets with a minimum of 20 million read pairs per sample; (2) diverse tissue representation for each species where available; and (3) taxonomic breadth across all eight molluscan classes. For species with multiple available datasets, preference was given to those with greater sequencing depth or tissue diversity. When multiple SRA runs existed for a single biological sample, reads were concatenated prior to assembly.

A concerted effort was made to include species from underrepresented classes to ensure broad taxonomic representation across all eight molluscan classes: Bivalvia, Caudofoveata, Cephalopoda, Gastropoda, Monoplacophora, Polyplacophora, Scaphopoda, and Solenogastres (Figure 1). The final dataset comprises 299 species with the following class distribution: Gastropoda (139 species, 46.5%), Bivalvia (61 species, 20.4%), Solenogastres (34 species, 11.4%), Cephalopoda (26 species, 8.7%), Polyplacophora (25 species, 8.4%), Caudofoveata (8 species, 2.7%), Scaphopoda (5 species, 1.7%), and Monoplacophora (1 species, 0.3%).

**Figure 1.**
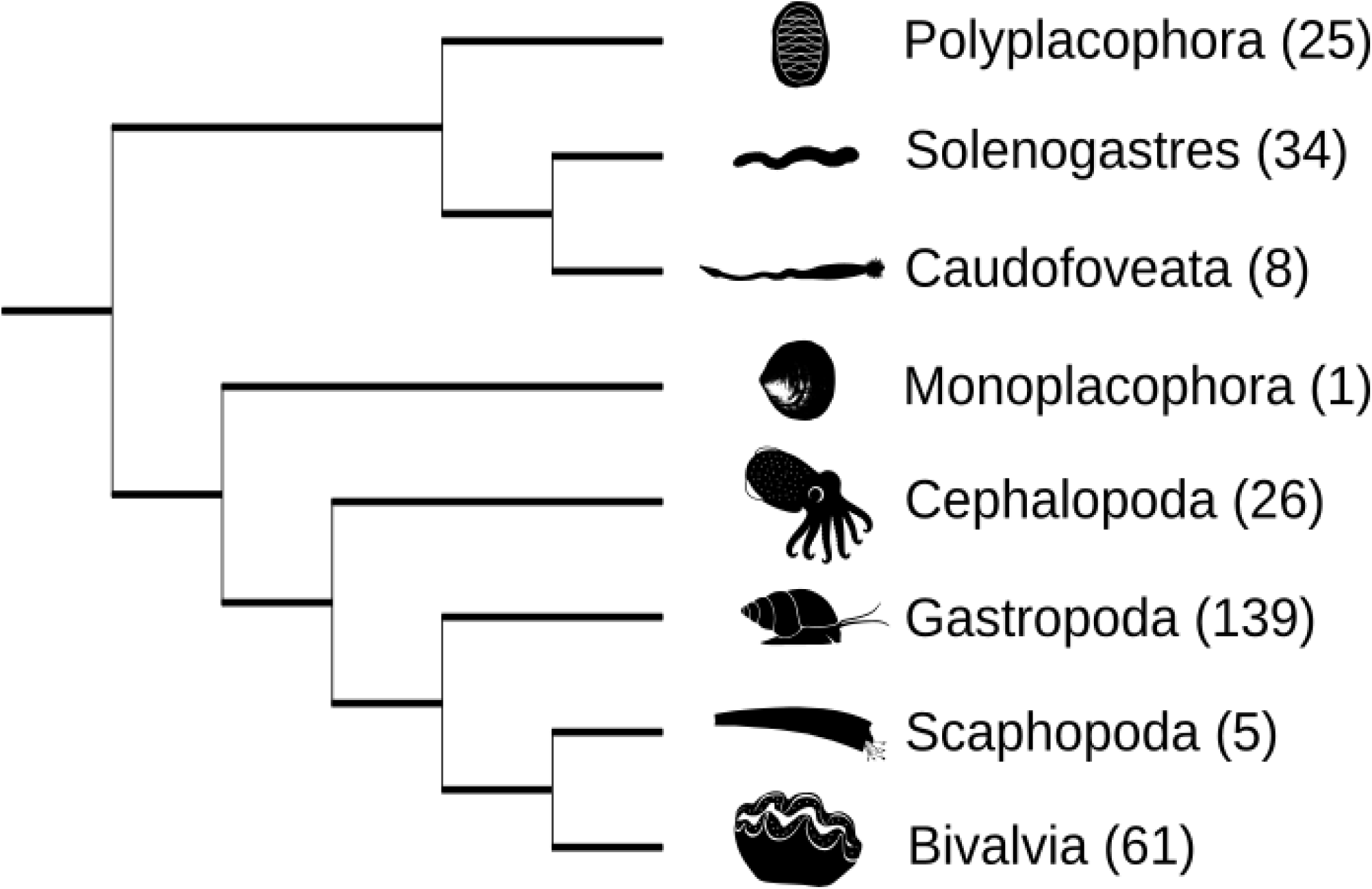
Taxonomic coverage of MolluscaGenes. Cladogram showing the distribution of 299 species across all eight recognized molluscan classes, species numbers for each clade in parenthesis.

In addition to raw reads, pre-assembled transcriptomes were sourced from legacy databases including MateDB (v1.0) and MolluscDB (v1.0). These pre-assembled datasets were processed through the EvidentialGene pipeline (Gilbert 2019) to reduce redundancy and select the highest-quality transcripts for inclusion, applying identical quality thresholds as for newly assembled transcriptomes. Datasets were excluded if they contained fewer than 10,000 predicted coding sequences after processing, or if BUSCO completeness scores fell below 50% against the Metazoa database (odb10). A complete summary of the included species, their corresponding database ID, tissue or developmental stage, SRA accession number, and data source is provided in Supplementary Table S1.

### 2.2 Transcriptome Assembly and Annotation

Raw reads were quality-controlled using fastp v0.23.4 (Chen et al. 2018) with the following parameters: adapter auto-detection enabled, minimum quality score of 20 for sliding window trimming (window size 4), minimum read length of 50 bp after trimming, and low-complexity filtering enabled (threshold 30%). Reads with more than 40% bases below quality 15 were discarded.

The cleaned reads were assembled *de novo* using the nf-core/denovotranscript pipeline v1.2.1 (Bhojwani et al. 2024), which employs a multi-assembler strategy to maximize transcript recovery. The pipeline executes Trinity v2.15.1 (Grabherr et al. 2011) with default parameters (minimum contig length 200 bp, k-mer size 25) and rnaSPAdes v3.15.5 (Bushmanova et al. 2019) with multiple k-mer sizes (k = 25, 49, 73). Assemblies from both tools were merged and processed through EvidentialGene v2023.07.15 (Gilbert 2019) for redundancy reduction. EvidentialGene identifies open reading frames (minimum length 100 amino acids), clusters transcripts at 98% identity, and selects optimal representatives based on coding potential and completeness, retaining both “main” (primary) and “alternate” (secondary) transcript models. Assembly quality was assessed using BUSCO v5.4.7 (Manni et al. 2021) against the Metazoa odb10 database (954 orthologs). Assemblies with BUSCO completeness below 50% were flagged for manual review; those below 30% were excluded from the database.

Protein sequences were functionally annotated using a multi-step approach. Primary annotations were assigned using DIAMOND v2.1.8 (Buchfink et al. 2015) blastp searches against the NCBI RefSeq protein database (release 220, September 2024) with an E-value threshold of 1e-20 and a maximum of 5 target sequences per query. Protein domains were identified using InterProScan v5.63-95.0 (Jones et al. 2014) with the following member databases: Pfam 35.0, SMART 9.0, CDD 3.20, SUPERFAMILY 1.75, and Gene3D 4.3.0.

Gene Ontology (GO) terms were assigned via InterProScan domain mappings and supplemented with eggNOG-mapper v2.1.12 (Cantalapiedra et al. 2021) using the eggNOG database v5.0. KEGG pathway assignments were obtained from eggNOG-mapper annotations.

### 2.3 Database Construction

The final assembled transcripts and translated protein sequences were formatted into searchable databases organized hierarchically by taxonomic class and species. The complete database comprises over 17 million transcript sequences (16.8 Gb total nucleotide sequence) and over 17 million predicted protein sequences (3.3 Gb total amino acid sequence), representing 299 molluscan species.

FASTA files are organized in three formats: (1) individual species files containing all transcripts for a given species; (2) class-level files combining all species within each molluscan class; and (3) a combined phylum-wide file for comprehensive searches. Each sequence header includes a standardized identifier encoding species, tissue source, and transcript classification (main vs. alternate). Searchable databases were constructed for the BLAST+ suite v2.14.0 using makeblastdb with the following parameters: nucleotide databases (blastn) with default settings, and protein databases (blastp/blastx) with taxonomic information embedded. DIAMOND databases v2.1.8 (Buchfink et al. 2015) were built using the --taxonmap and --taxonnodes options to enable taxonomic filtering of search results.

Command-line access is facilitated through local execution of these search tools, with wrapper scripts provided for common query types. The databases are also provided as downloadable FASTA files for use with custom tools and scripts. Example scripts, detailed tutorials, and a quick-start guide for performing queries are provided in the project’s official GitHub repository (https://github.com/invertome/molluscagenes). The repository includes workflows for batch processing of query sequences and integration with downstream phylogenetic analysis pipelines. The overall database construction pipeline is illustrated in Figure 2.

**Figure 2.**
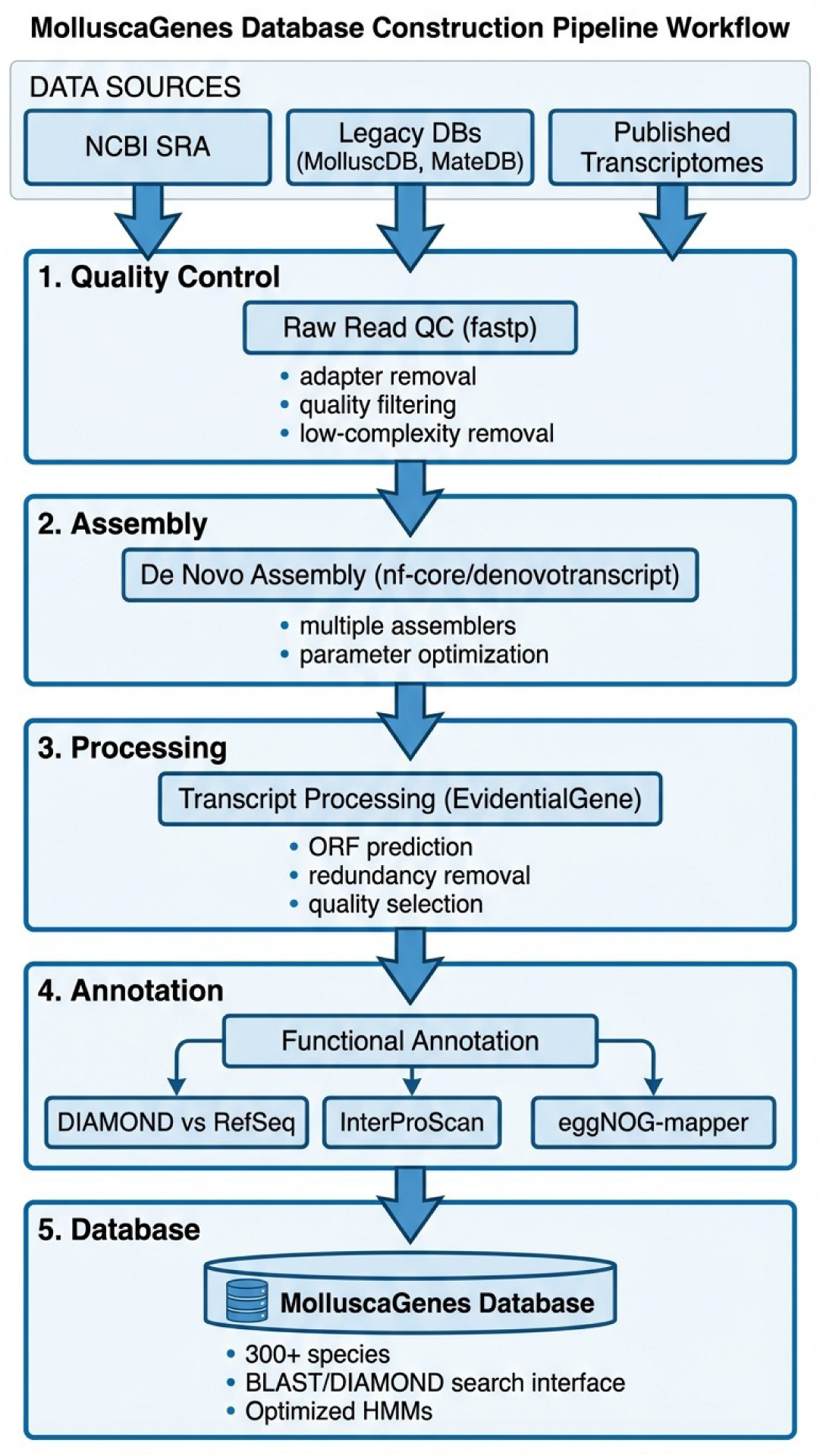
MolluscaGenes database construction pipeline. Schematic overview of the data acquisition, processing, and database construction workflow. Raw sequence data from NCBI SRA and pre-assembled transcriptomes from legacy databases are processed through quality control (fastp), de novo assembly (nf-core/denovotranscript), and redundancy reduction (EvidentialGene) to generate optimized protein-coding sequences. The final database supports BLAST, DIAMOND, and HMM-based searches.

Database updates will be issued on a regular basis to incorporate newly published molluscan transcriptomes, with version numbering following semantic versioning conventions. Users can subscribe to release notifications via the GitHub repository.

### 2.4 Development of Optimized HMMs

To facilitate sensitive, deep-homology searches across Mollusca, we developed a suite of 196 optimized Hidden Markov Models (HMMs) encompassing Pfam domain families distributed across 50 functional categories of fundamental relevance to molluscan biology, including innate immunity, developmental signaling, calcium signaling, voltage-gated ion channels, G protein-coupled receptors, ligand-gated ion channels, shell biomineralization, epigenetic regulation, and additional categories spanning chemoreception, circadian rhythms, neuropeptide processing, toxin biology, and synaptic function. Initial HMMs were sourced from Pfam 35.0 for domain families listed in Supplementary Table S2.

These generic models were optimized for molluscan-specific sequence diversity using the TIAMMAt workflow v1.0 (Tassia et al. 2021). The iterative optimization process proceeds as follows: (1) the initial Pfam HMM is used to search MolluscaGenes with hmmsearch (HMMER v3.3.2) at a permissive E-value threshold of 0.001; (2) recovered sequences are aligned using MAFFT v7.520 with the E-INS-i algorithm; (3) a new HMM is built from this alignment using hmmbuild with default parameters; (4) the refined HMM is used to search MolluscaGenes again, recovering additional divergent sequences; (5) steps 2--4 are iterated until no new sequences are recovered (typically 3--5 iterations). The final optimized HMM incorporates molluscan-specific sequence variation absent from the original vertebrate- and ecdysozoan-biased Pfam models.

Validation of optimized HMMs was performed by comparing detection rates against the original Pfam models on six held-out molluscan proteomes (*Aplysia californica*, *Crassostrea gigas*, *Lingula anatina*, *Lottia gigantea*, *Octopus bimaculoides*, and *Pomacea canaliculata*). False positive rates were assessed by searching shuffled sequence databases. All 196 optimized HMMs are provided in the MolluscaGenes repository in HMMER3 format, with accompanying documentation describing the gene families targeted and recommended E-value thresholds for different search stringencies.

### 2.5 Phylum-Wide Characterization of the nAChR Superfamily

To characterize the diversity and evolutionary history of the nAChR superfamily across Mollusca, including the recently described chemotactile receptors (CRs) (Giesen et al. 2020; Allard et al. 2023), we employed a comprehensive homology search strategy against the MolluscaGenes protein database. A curated reference set of 191 nAChR and CR protein sequences was assembled from three molluscan species (*Aplysia californica*, *Lymnaea stagnalis*, *Octopus bimaculoides*) and two outgroup taxa (*Drosophila melanogaster*, *Homo sapiens*), encompassing characterized CR sequences from *O. bimaculoides* (Supplementary Materials S3). NCBI BLAST+ blastp searches were conducted with a permissive E-value threshold of 1e-5 and a maximum of 5,000 target sequences per query to ensure the recovery of divergent superfamily members.

Candidate sequences were subjected to stringent multi-criterion filtering: E-value $\leq$ 1e-10, sequence identity $\geq$ 30%, query coverage $\geq$ 40%, minimum subject length $ \geq$ 300 amino acids, and minimum bitscore $\geq$ 100. Deduplication retained only the highest-scoring hit per subject sequence, yielding 4,057 unique candidates from the initial filtering stage. These candidates were subsequently screened with our molluscan-optimized HMM for the neurotransmitter-gated ion channel ligand-binding domain (Neur_chan_LBD; Pfam: PF02931) via hmmsearch (HMMER v3.3.2) (Eddy 2011). Validation of characteristic Cys-loop receptor domains was performed through InterProScan annotation (Jones et al. 2014).

Multiple sequence alignment of the confirmed candidates was generated using MAFFT v7.526 (Katoh and Standley 2013) with the E-INS-i algorithm (`--genafpair --maxiterate 1000`), selected for its suitability for sequences containing conserved motifs (the Cys-loop and transmembrane domains) separated by variable-length regions. The resulting alignment (3,586 sequences, 4,176 columns) was trimmed with ClipKIT v2.3.0 (Steenwyk et al. 2020) under the kpic-smart mode, which retains parsimony-informative and constant sites while removing ambiguously aligned positions.

Maximum-likelihood phylogenetic inference was conducted with IQ-TREE2 v2.4.0 (Nguyen et al. 2015) under the Q.pfam+G8 substitution model (a Pfam-specific rate matrix with an eight-category discrete gamma distribution). Branch support was assessed via 1,000 ultrafast bootstrap replicates (`-B 1000`) with near-zero branch length optimization (`-bnni`) and the approximate Bayes test (`-abayes`). The `--safe` flag was enabled to accommodate the computational demands of this large-scale alignment. The tree was rooted with GABA-A and glycine receptor sequences from representative metazoans as outgroups (Supplementary Materials S3).

Clade assignment employed a reference-guided hierarchical approach. TreeCluster (threshold 5.0, minimum support 0.90) partitioned the phylogeny into 180 clusters and 48 singletons, after which the 191 pre-assigned reference sequences served as anchors for majority-vote clade designation within reference-containing clusters. Sequences in clusters lacking reference representatives were assigned via nearest-neighbor classification. This procedure delineated 15 distinct clades within the nAChR superfamily across the phylum.

Functional annotation of the identified superfamily members encompassed the extraction of ion selectivity determinants from the TM2 region (GEK, GER, PAR, and TEK motifs), assessment of Cys-loop motif conservation (YXCC), signal peptide prediction, and glycosylation site detection. Rogue taxon identification, taxonomic subsampling procedures, and the motif extraction pipeline are described in detail in the Supplementary Methods.

## 3. Results

### 3.1 Overview of the MolluscaGenes Resource

The MolluscaGenes database consolidates transcriptomic data from 299 species representing all eight recognized classes of Mollusca. The taxonomic breadth of MolluscaGenes is illustrated in Figure 1, which shows the distribution of included species across the molluscan phylogeny. Gastropoda constitutes the most heavily represented class (139 species, 46.5%), reflecting both taxonomic diversity and data availability, followed by Bivalvia (61 species, 20.4%) and Solenogastres (34 species, 11.4%). Cephalopoda (26 species), Polyplacophora (25 species), Caudofoveata (8 species), Scaphopoda (5 species), and Monoplacophora (1 species) provide representation of the remaining classes, ensuring comprehensive coverage for comparative evolutionary analyses across the phylum.

### 3.2 Database Functionality and User Interface

MolluscaGenes is accessible via a set of downloadable files with command-line instructions, available online at: https://invertome.github.io/molluscagenes/. Three primary query pathways are supported: (1) sequence homology searches via BLAST or DIAMOND for identifying similar sequences across the database; (2) HMM-based searches for sensitive detection of protein family members, particularly effective for divergent homologs; and (3) bulk taxonomic downloads for obtaining complete transcriptome datasets for specific species or clades.

### 3.3 Molluscan-Optimized HMMs

Application of the TIAMMAt iterative optimization workflow to 196 Pfam seed alignments yielded 196 molluscan-optimized HMMs with substantially increased sensitivity for the detection of divergent homologs across MolluscaGenes. Validation against six held-out molluscan proteomes demonstrated that the optimized HMMs recovered 20.4% more sequences on average relative to the original Pfam models at equivalent E-value stringency, with the most pronounced gains observed for those in the “Developmental signaling” category (+1369.1%). False positive assessment against shuffled sequence databases confirmed that the optimized models retain high specificity (false positive rate $<$ XXXX%). This augmented sensitivity proved particularly valuable for the identification of highly divergent superfamily members that elude conventional BLAST-based searches, as demonstrated in the nAChR case study below.

### 3.4 Phylum-Wide Characterization of the nAChR Superfamily

#### 3.4.1 Identification and Clade Diversity

The application of molluscan-optimized HMMs and reference-guided phylogenetic classification to MolluscaGenes yielded a total of 3,586 nAChR superfamily sequences from more than 190 species spanning all eight recognized molluscan classes, along with outgroup taxa from Nematoda (*Caenorhabditis elegans*), Insecta (*Drosophila melanogaster*), and Vertebrata (*Homo sapiens*). Maximum-likelihood phylogenetic inference (IQ-TREE2, Q.pfam+G8 model, 1,000 ultrafast bootstrap replicates with aBayes support) resolved these sequences into 15 well-supported monophyletic clades (Figure 3). The five largest clades, nAChR-L (795 sequences), AChRB (457), Alpha-A10a (374), Alpha-A2 (370), and Alpha-L (278), collectively encompassed approximately 63% of the dataset and exhibited broad taxonomic conservation across multiple molluscan classes. The remaining ten clades ranged from moderately sized assemblages such as the Cys-less (231 sequences) and Non-alpha-1 (221) clades to smaller, more taxonomically restricted lineages including Alpha-6/7 (61 sequences). Notably, the CR (94 sequences) and CR-like (97 sequences) clades, previously characterized primarily in cephalopods, emerged as phylogenetically distinct lineages within this broader superfamily framework. A total of 125 sequences (3.5% of the dataset) were flagged as rogue taxa, the majority originating from the nAChR-L (63 sequences) and Alpha-L (34 sequences) clades, and were excluded from downstream clade-level analyses.

**Figure 3.**
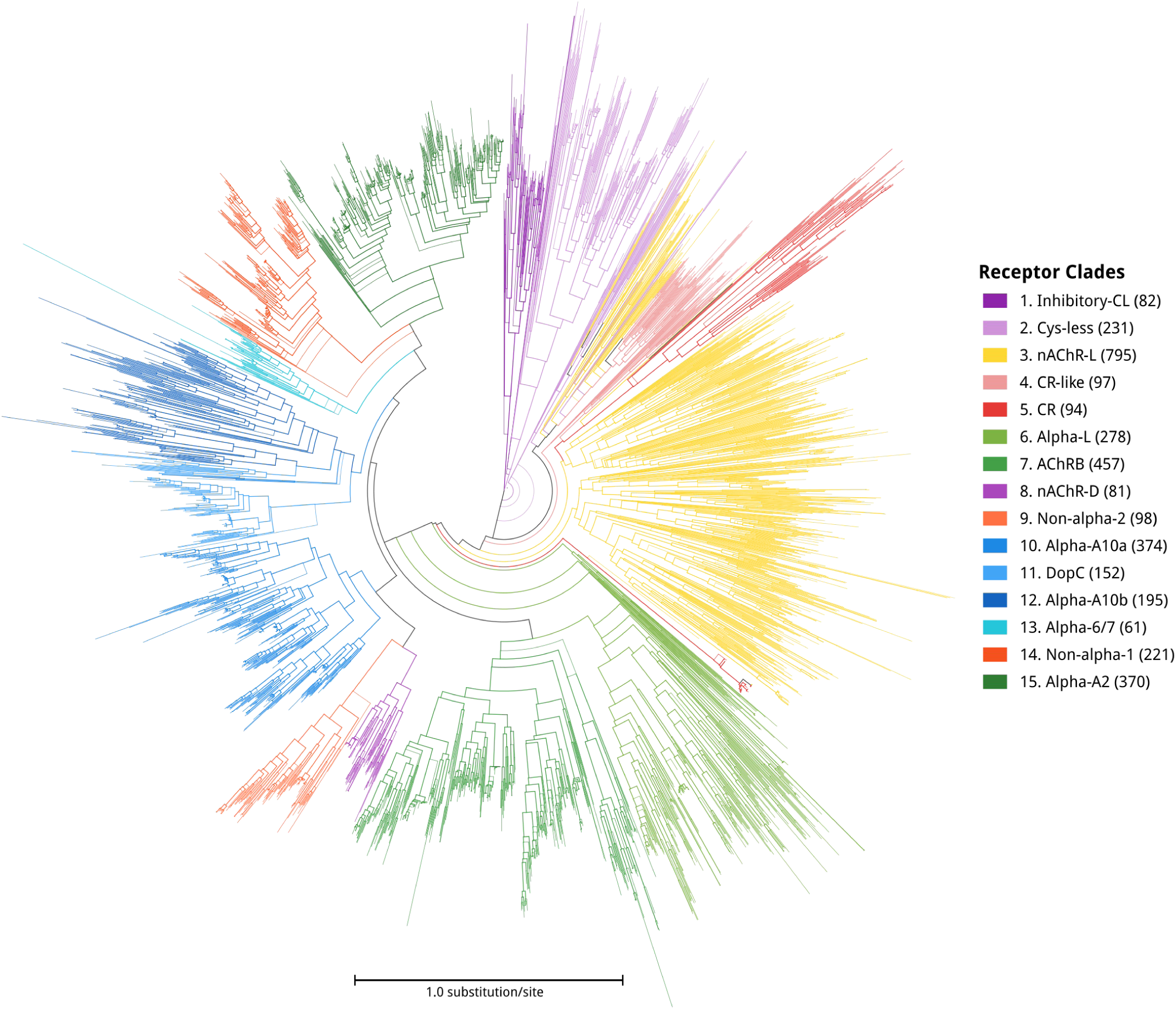
Phylum-wide phylogeny of the molluscan nAChR superfamily. Maximum-likelihood phylogenetic tree of 3,586 nAChR superfamily sequences identified across 190+ molluscan species. Branches are colored by clade assignment (15 clades identified via TreeCluster with reference-guided mapping). The tree was inferred using IQ-TREE2 with the Q.pfam+G8 substitution model and 1,000 ultrafast bootstrap replicates. Outgroup sequences (GABA-A and glycine receptors) are shown in gray. Scale bar indicates substitutions per site.

#### 3.4.2 Taxonomic Distribution and Lineage-Specific Expansions

The distribution of nAChR superfamily clades across molluscan classes revealed pronounced patterns of lineage-specific enrichment (Figure 4; Supplementary Figures S3, S6).

**Figure 4.**
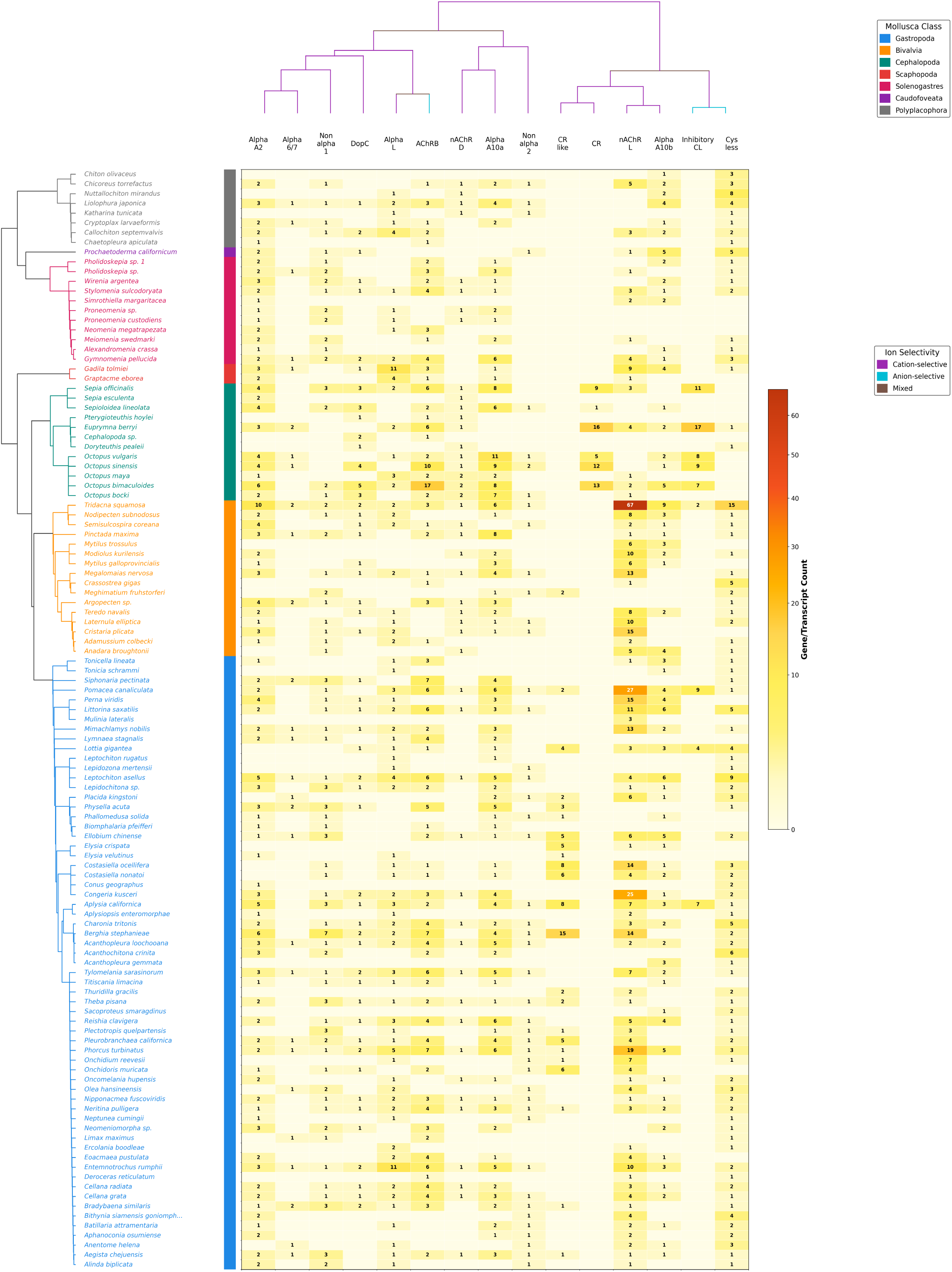
Distribution of nAChR superfamily clades across molluscan species. Heatmap showing the number of sequences assigned to each of the 15 nAChR clades for each species in the analysis. Species are ordered by taxonomic class, with class-level annotations shown on the left axis. Clade names are shown along the top axis with color coding matching Figure 3. The CR and CR-like clades show broad taxonomic distribution, with enrichment in cephalopod species.

Gastropoda, the most densely sampled class (approximately 78 species), contributed the largest proportion of sequences and dominated the nAChR-L and Alpha-A2 clades, consistent with the substantial transcriptomic representation of this class within MolluscaGenes.

Bivalvia (approximately 21 species) exhibited conspicuous enrichment in the Alpha-A10a, Alpha-A10b, and AChRB clades, a pattern that appears concordant with previous reports of extensive nAChR gene family expansion in bivalve genomes (Li et al. 2019). Cephalopoda (approximately 19 species), despite more limited species sampling, displayed marked enrichment in the Alpha-A10a, DopC, CR, and CR-like clades, the latter two of which suggest a distinctive trajectory of receptor diversification in this lineage. Representation from Polyplacophora (approximately 6 species) spanned multiple clades at lower sequence counts, whereas Scaphopoda, Solenogastres, and Caudofoveata contributed sparse but phylogenetically informative sequences. These distributional asymmetries may reflect genuine differences in gene family expansion across molluscan lineages, although the influence of differential transcriptomic sampling depth and tissue-specific expression on the observed patterns warrants consideration.

#### 3.4.3 Domain Architecture and Sequence Conservation

The canonical Cys-loop receptor architecture, comprising an extracellular ligand-binding domain (LBD) and four transmembrane helices (TM1--TM4), was conserved across the majority of identified sequences (Dent 2006; Jones and Sattelle 2007). A notable exception was observed in the Cys-less clade (231 sequences) and the Inhibitory-CL clade (82 sequences), both of which lacked the characteristic YXCC Cys-loop motif that is otherwise broadly conserved across the superfamily. Analysis of ion selectivity determinants in the TM2 region revealed that most clades exhibited predominantly cation-selective signatures (GEK-type motifs), consistent with classical excitatory nAChR function (Supplementary Figures S2, S5). The CR clade, by contrast, displayed mixed ion selectivity patterns, whereas the Inhibitory-CL and Cys-less clades harbored PAR-type and other anion-selective motifs suggestive of inhibitory channel properties (Supplementary Figure S7). Preliminary assessment of the ligand-binding domain suggests that several divergent clades may harbor substitutions at positions associated with agonist coordination; however, the functional implications of these modifications remain to be determined through experimental validation.

#### 3.4.4 Chemotactile Receptors and CR-like Sequences

Phylogenetic analysis resolved two distinct clades related to the chemotactile receptors (CRs) first described in octopods (Giesen et al. 2020): the CR clade (94 sequences) and the CR-like clade (97 sequences), which together constituted 5.3% of the dataset (191 sequences). The CR clade was predominantly composed of cephalopod sequences, with the highest representation observed in *Euprymna berberorum* (21 sequences), *Octopus bimaculoides* (15), *Octopus sinensis* (14), and *Sepia officinalis* (14), consistent with the cephalopod-enriched expansion previously documented (Allard et al. 2023). The CR-like clade, by contrast, exhibited a substantially broader taxonomic distribution that extended across gastropods, bivalves, and polyplacophorans, suggesting that CR-related sequences are not restricted to cephalopods but rather represent a more ancient component of the molluscan nAChR repertoire.

The intronless genomic structure characteristic of cephalopod CRs, which has been attributed to a retroposition origin in *O. bimaculoides* (Giesen et al. 2020), remains to be verified across additional cephalopod genomes. Examination of ligand-binding domain residues revealed that the CR clade harbors modifications at positions typically associated with acetylcholine coordination, paralleling analogous substitutions observed in certain divergent clades (e.g., Cys-less, nAChR-D). This convergence of binding pocket modifications across phylogenetically distant lineages raises the possibility of independent evolutionary transitions toward alternative ligand specificities within the molluscan nAChR superfamily, although functional characterization will be required to substantiate this hypothesis.

## 4. Discussion

### 4.1 MolluscaGenes as a Foundational Resource

The fragmentation of molluscan transcriptomic data across disparate repositories has constituted a persistent impediment to comparative analyses spanning the breadth of this extraordinarily diverse phylum. Existing resources, including Ensembl Metazoa and OrthoDB (Howe et al. 2021; Waterhouse et al. 2013), provide expansive multi-phylum coverage; however, this taxonomic breadth necessarily sacrifices the dense within-lineage sampling required for fine-grained evolutionary reconstruction. MolluscaGenes addresses this limitation by consolidating transcriptomic data from 299 species representing all eight recognized molluscan classes, thereby enabling phylogenetically informed analyses at a resolution previously unattainable within a single resource. The combination of curated ortholog sets, molluscan-optimized HMMs, and standardized annotation pipelines provides a foundation for investigations ranging from deep phylogenomics to the characterization of lineage-specific molecular innovations.

The utility of MolluscaGenes for evolutionary biology is exemplified by the study of biomineralization, one of the most ancient and well-characterized phenomena in the phylum. Molluscan shells represent a paradigmatic system for understanding the molecular mechanisms underlying biologically controlled mineralization (Kocot et al. 2016). The mantle secretome, which governs the formation of the organic matrix directing shell construction, exhibits remarkable evolutionary plasticity; shell matrix proteins (SMPs) display evidence of both deeply conserved gene families and lineage-specific expansions that contribute to the extraordinary diversity of shell architectures observed across the Mollusca (Jackson et al. 2006; McDougall et al. 2013). The elucidation of SMP evolutionary trajectories requires dense taxonomic sampling of the kind MolluscaGenes provides, as demonstrated by genomic analyses of *Crassostrea gigas* that revealed both conserved and novel components of the biomineralization toolkit (Zhang et al. 2012; Takeuchi et al. 2012). By enabling phylum-wide comparisons of mantle-expressed gene families, MolluscaGenes facilitates the reconstruction of ancestral biomineralization programs and the identification of lineage-specific innovations.

The resource holds equally fundamental significance for molluscan neurobiology. Within the Cephalopoda, organisms such as octopuses, squids, and cuttlefish possess nervous systems of extraordinary complexity, with *Octopus vulgaris* harboring approximately 500 million neurons, a quantity comparable to that observed in small primates and representing a striking instance of convergent evolution with vertebrates (Young 1963; Budelmann 1995; Shigeno et al. 2018). The sequencing of the *Octopus bimaculoides* genome revealed surprising molecular innovations, including massive expansions in protocadherin gene families (168 genes, compared to 17--25 in other invertebrates) and C2H2 zinc-finger transcription factors, gene families previously considered uniquely expanded in vertebrates (Albertin et al. 2015). These discoveries suggest that complex neural organization may arise through convergent molecular mechanisms across distantly related lineages (Schnell and Clayton 2021; Moroz 2021). MolluscaGenes enables systematic comparative transcriptomic analyses between cephalopod and gastropod nervous systems, facilitating the identification of conserved synaptic scaffolding proteins, neural adhesion molecules, and lineage-specific innovations that underpin the independent elaboration of neural complexity across molluscan classes.

Beyond fundamental research, MolluscaGenes provides substantial opportunities for applied science. The phylum Mollusca constitutes a prolific source of bioactive compounds, exemplified by the cone snail-derived conotoxins that have yielded the analgesic ziconotide and continue to furnish promising pharmacological leads (Lewis et al. 2012; Gao et al. 2017). Dense taxonomic sampling across venomous and non-venomous lineages facilitates bioprospecting through the identification of novel toxin gene families and their evolutionary diversification. The resource additionally supports the nascent field of cellular agriculture: the development of cultured seafood from molluscan cell lines requires comprehensive molecular characterization of growth factors, extracellular matrix components, and differentiation pathways (Rubio et al. 2020). Furthermore, the identification of immune-related gene families across bivalve species provides a foundation for understanding disease resistance mechanisms in commercially important species and for developing molecular markers for selective breeding programs in aquaculture (Song et al. 2015).

### 4.2 Evolutionary Diversity of the Molluscan nAChR Superfamily

The phylum-wide characterization of nAChR superfamily members presented here constitutes the most comprehensive evolutionary framework for this receptor family in the Mollusca to date. The delineation of 15 distinct clades corroborates and substantially extends the classification proposed by Li et al. (2019), who identified major nAChR subfamilies from a more limited sampling of bivalve genomes. In vertebrates, the diversification of the nAChR superfamily has been shaped predominantly by two rounds of whole-genome duplication (WGD) that generated paralogous receptor subunits subsequently refined by subfunctionalization and gene loss (Pedersen et al. 2019). The molluscan nAChR repertoire, by contrast, appears to have expanded primarily through tandem duplication and extensive sequence divergence, a mechanism that may confer greater evolutionary flexibility for the generation of novel receptor pharmacologies within individual lineages.

The identification of divergent nAChR clades harboring substitutions at canonical ligand-binding residues carries substantial implications for the understanding of molluscan sensory ecology. The convergence of binding pocket modifications across phylogenetically distant clades warrants investigation as a potential indicator of independent evolutionary transitions toward the detection of non-canonical ligands. Gastropod chemosensory systems, which employ specialized structures such as rhinophores, osphradia, and oral tentacles for the detection of waterborne chemical stimuli (Croll 1983), represent particularly compelling candidates for the expression of such divergent receptors. Recent characterization of G protein-coupled receptors (GPCRs) in the rhinophore epithelia of opisthobranch gastropods (Cummins et al. 2009), together with documentation of extensive GPCR expansions across lophotrochozoan lineages (Nath and Bhattacharya 2025), suggests that the peripheral chemosensory repertoire of gastropods may be considerably more complex than previously appreciated. The possibility that divergent nAChR clades contribute an ionotropic component to these chemosensory pathways merits systematic functional exploration.

The placement of chemotactile receptors (CRs) within the broader nAChR superfamily framework illuminates the evolutionary origins of this cephalopod-enriched receptor class. The intronless genomic structure characteristic of CRs is consistent with an origin via retroposition from an intron-containing nAChR ancestor (Kaessmann et al. 2009; Vinckenbosch et al. 2006), a mechanism that may have facilitated rapid functional divergence following retrotransposition. Cryo-electron microscopy structures reveal that while the overall CR channel architecture remains conserved with ancestral nAChRs, the ligand-binding pocket has undergone substantial remodeling toward an exceptionally hydrophobic configuration, enabling the detection of water-insoluble compounds such as terpenoids (Allard et al. 2023). This structural adaptation appears well suited to the distributed nervous system of octopus arms, in which approximately two-thirds of the animal’s neurons reside and function semi-autonomously in the exploration and chemotactile assessment of the environment (Levy et al. 2015; Zullo and Hochner 2019). By situating CRs within their broader phylogenetic context, MolluscaGenes enables the identification of shared and derived features that distinguish this lineage from canonical nAChR clades.

The evolutionary plasticity of the Cys-loop receptor superfamily, of which nAChRs constitute a major division, further contextualizes the diversity observed in the present study. Previous work has demonstrated that changes in ion selectivity, from cation-selective to anion-selective channels, can be accomplished through remarkably few mutations in the pore-lining TM2 domain (Pirri et al. 2015; Jones and Sattelle 2008). The identification of anion-selective nAChR-related channels derived from cation-selective ancestors in multiple invertebrate lineages (Nierop et al. 2005) underscores the evolutionary lability of this functional property. The phylogenetic framework established here provides the taxonomic resolution necessary for testing whether multiple independent transitions in ion selectivity have occurred within the molluscan nAChR superfamily, and for reconstructing the ancestral states from which such transitions may have arisen.

The detection of CR-like sequences across non-cephalopod molluscan classes represents a notable finding from an evolutionary classification perspective. Whether these sequences represent functional homologs of cephalopod CRs, ancestral forms that retain canonical neurotransmitter-gated function, or pseudogenes undergoing neutral evolution constitutes a compelling question that the present phylogenetic analysis alone cannot resolve. The broad taxonomic distribution of CR-like sequences across gastropods, bivalves, and polyplacophorans suggests that the ancestral gene from which cephalopod CRs arose was present in the molluscan common ancestor, with subsequent neofunctionalization occurring specifically within the coleoid lineage. Functional characterization of CR-like sequences from non-cephalopod species through heterologous expression and electrophysiology will be essential for distinguishing among these evolutionary scenarios.

In aggregate, the nAChR superfamily analysis presented here demonstrates the capacity of MolluscaGenes to enable the identification of novel receptor diversity, the reconstruction of gene family evolutionary history across an entire phylum, and the formulation of testable hypotheses regarding receptor evolution and function. The scope of diversity revealed, with 15 clades encompassing more than 3,500 sequences from over 190 species, exceeds the 29 nAChR subunits characterized in the extensively studied model organism *Caenorhabditis elegans* (Jones and Sattelle 2008) and underscores the magnitude of unexplored receptor diversity within the Mollusca.

### 4.3 Future Directions

MolluscaGenes is designed as a living resource, with planned annual updates incorporating newly deposited transcriptomic datasets, expanded taxonomic coverage for underrepresented classes (particularly Monoplacophora and Caudofoveata), and the integration of tissue-specific and developmental expression data. The incorporation of quantitative expression profiles will enable the transition from sequence-level annotation to spatiotemporal characterization of gene expression across molluscan body plans.

The rapid maturation of single-cell RNA sequencing (scRNA-seq) technologies in molluscan systems constitutes a particularly promising frontier for the resource. Recent atlases of the *Octopus vulgaris* brain have begun to delineate the cellular diversity of cephalopod neural tissue at unprecedented resolution (Styfhals et al. 2022), while complementary efforts in squid (Duruz et al. 2023) and gastropod ganglia (Ramirez et al. 2024) are establishing cell-type inventories across divergent molluscan lineages. MolluscaGenes provides a comprehensive reference framework against which these emerging single-cell datasets can be annotated, enabling the identification of cell-type-specific receptor expression patterns and the comparative analysis of neural cell populations across the phylum.

The application of deep learning approaches to the prediction of protein function from sequence constitutes an additional avenue of substantial promise. Methods such as DeepFRI and ProteInfer (Gligorijevic et al. 2021; Sanderson et al. 2023) demonstrate that neural networks trained on large, diverse protein datasets can predict Gene Ontology terms and functional annotations with considerable accuracy. The breadth of divergent sequences cataloged in MolluscaGenes, particularly within receptor superfamilies, provides training data that may improve the prediction of function for orphan sequences that lack close homologs in model organism databases.

Community engagement represents a fundamental component of the resource’s long-term sustainability. The MolluscaGenes GitHub repository provides a platform for user-contributed annotations, error reporting, and feature requests, fostering a collaborative model of database development. Expansion of user-driven curation, particularly for species-specific functional annotations and literature-linked gene records, will enhance the utility of the resource for the broader molluscan research community.

Several limitations inherent to transcriptome-based resources warrant explicit acknowledgment. The absence of a transcript from the database does not constitute evidence for the absence of the corresponding gene; tissue-restricted expression, developmental stage specificity, and transcript instability may all contribute to false negatives. Furthermore, transcriptomic data alone cannot resolve questions of genomic organization, including synteny, non-coding regulatory elements, and the distinction between tandem duplications and assembly artifacts. The future integration of chromosome-level genome assemblies, as they become available for additional molluscan species, will complement the transcriptomic foundation of MolluscaGenes and address these limitations.

The functional characterization of divergent nAChR clades identified in the present study represents a high-priority objective for subsequent investigation. *In situ* hybridization in gastropod chemosensory organs (rhinophores, oral tentacles, etc.) would establish the spatial expression patterns of divergent receptor subunits, while heterologous expression in *Xenopus* oocytes or mammalian cell lines would enable pharmacological profiling and ion selectivity determination. Structural modeling informed by the CRT1 cryo-EM structure (Allard et al. 2023) may further guide predictions regarding ligand binding and channel gating properties in these uncharacterized receptors.

## 5. Conclusion

In this study, we introduced MolluscaGenes, a comprehensive and taxonomically diverse transcriptomic resource encompassing 299 species across all eight recognized molluscan classes, accompanied by 196 molluscan-optimized HMMs for sensitive protein family identification. By providing a centralized platform with advanced search and annotation tools, MolluscaGenes addresses a fundamental limitation in molluscan genomics and facilitates the investigation of long-standing questions in evolutionary biology, neuroscience, and biotechnology.

The phylum-wide characterization of the nAChR superfamily, recovering 3,586 sequences distributed across 15 phylogenetic clades, demonstrates the capacity of this resource to unveil previously unrecognized receptor diversity at an unprecedented taxonomic scale. The resolution of chemotactile receptors and CR-like sequences as evolutionary clades within the nAChR superfamily, rather than as an independent gene family, provides a unified phylogenetic framework that recontextualizes the origin of these predominantly cephalopod receptors. The identification of substantial lineage-specific expansions, novel clades with substitutions in canonical ligand-binding residues, and nAChR paralogs within individual species collectively generate testable hypotheses regarding the molecular basis of receptor neofunctionalization across Mollusca.

MolluscaGenes constitutes a foundational resource that will accelerate continued discovery in molluscan biology, from comparative genomics and neurobiology to the elucidation of sensory system evolution.

## Author Contributions

JLP-M conceived and designed the study, developed the database and analysis pipelines, performed the phylogenetic analyses, and wrote the manuscript. PSK supervised the project and contributed to manuscript preparation and revision. All authors approved the final version.

## Acknowledgments

We thank Dr. Colleen Lawless for contributions to early stages of the chemotactile receptor analysis. Computational analyses were performed using the Unity HPC and AI platform, a multi-institutional high-performance computing resource managed by the University of Massachusetts Amherst and housed at the Massachusetts Green High Performance Computing Center (MGHPCC) in Holyoke, MA.

## Supplemental Data

Supplementary Table S1. contains the complete list of species included in MolluscaGenes, with corresponding database IDs, tissue sources, SRA accession numbers, and data sources.

Supplementary Table S3. the complete list and metadata information of the HMM profiles that were optimized for molluscs using the TIAMMAT pipeline.

Supplementary Table S3. contains the complete list of reference sequences used in the nAChR superfamily analysis and associated metadata.

## Data Availability Statement

The MolluscaGenes database is freely available at https://invertome.github.io/molluscagenes/. All sequences, optimized HMMs, and analysis scripts are deposited in the project GitHub repository at https://github.com/invertome/molluscagenes.

