## Supplementary Materials for "MolluscaGenes: A Transcriptomic Database for the Mollusca"

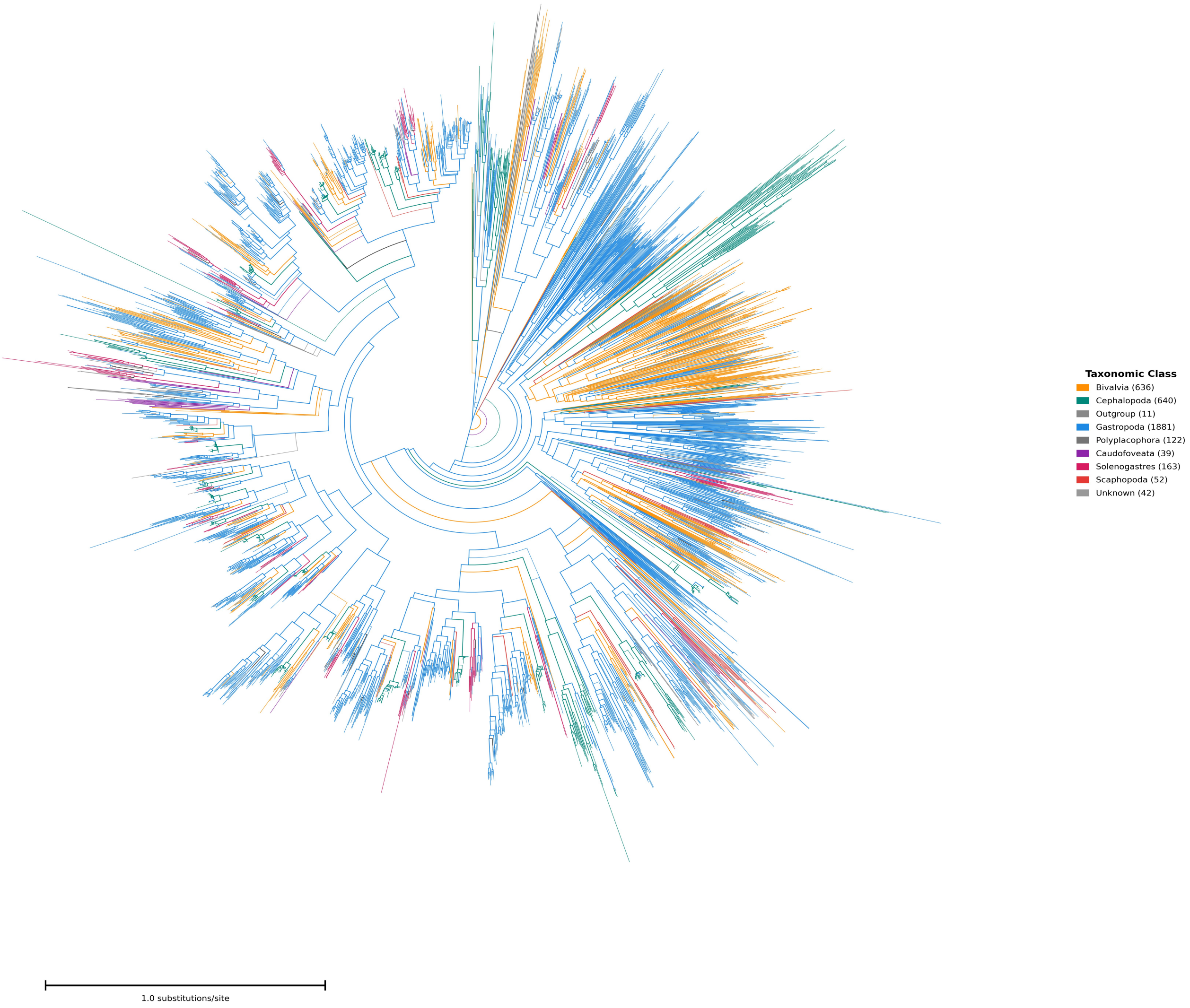


**Figure S1.** Phylogeny of the molluscan nAChR superfamily colored by taxonomic class. The same maximum-likelihood tree as Figure 3 (main text) with branches colored by molluscan class instead of clade assignment. Gastropoda (blue) constitutes the majority of sequences, followed by Bivalvia (orange) and Cephalopoda (teal). Outgroup sequences from Nematoda, Insecta, and Vertebrata are shown in gray.

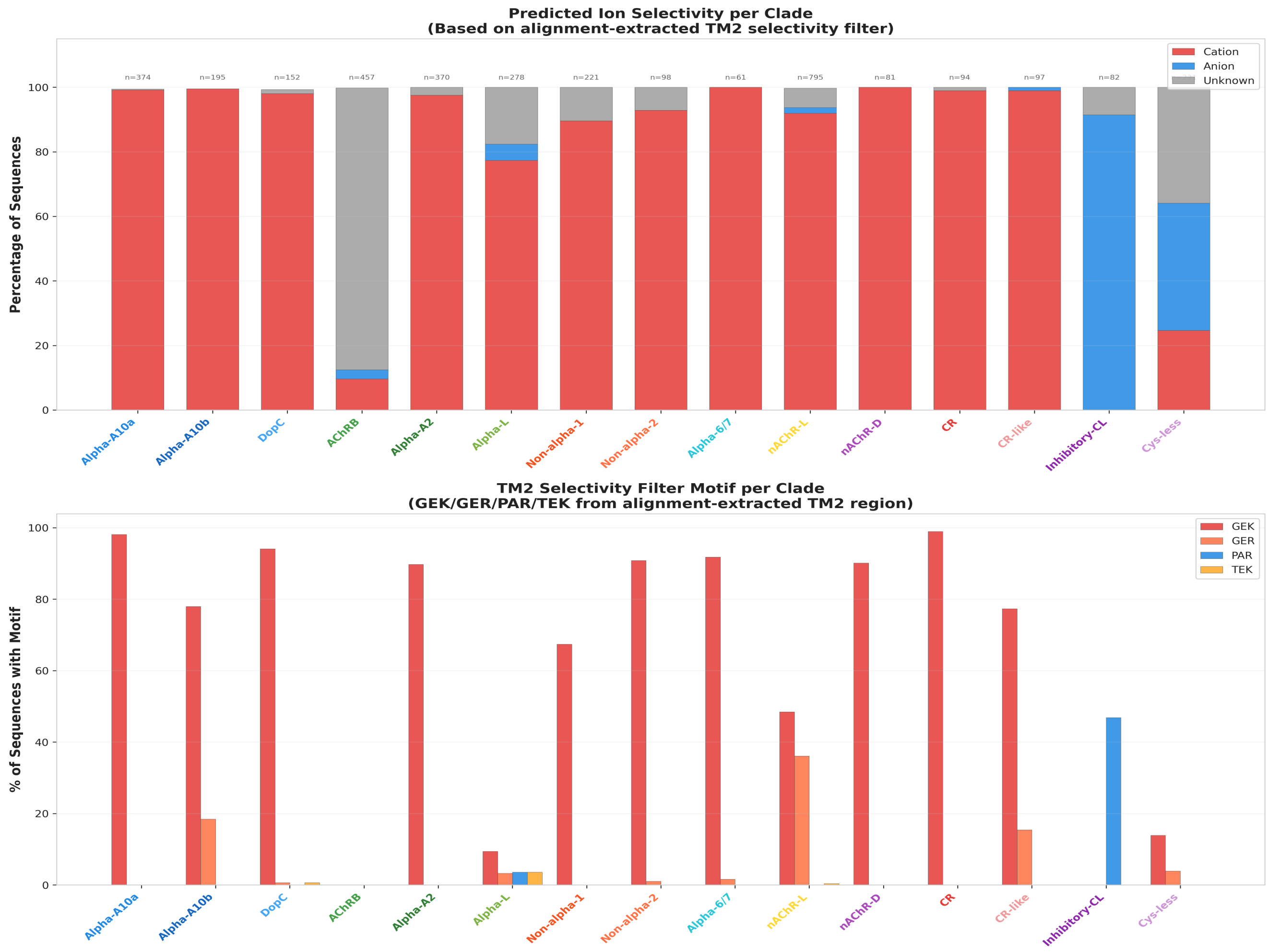


**Figure S2.** Predicted ion selectivity across nAChR superfamily clades. (Top) Proportion of sequences in each clade predicted to be cation-selective, anion-selective, or of undetermined selectivity, based on analysis of the TM2 selectivity filter region. (Bottom) Distribution of TM2 selectivity filter motifs (GEK, GER, PAR, TEK) across clades.

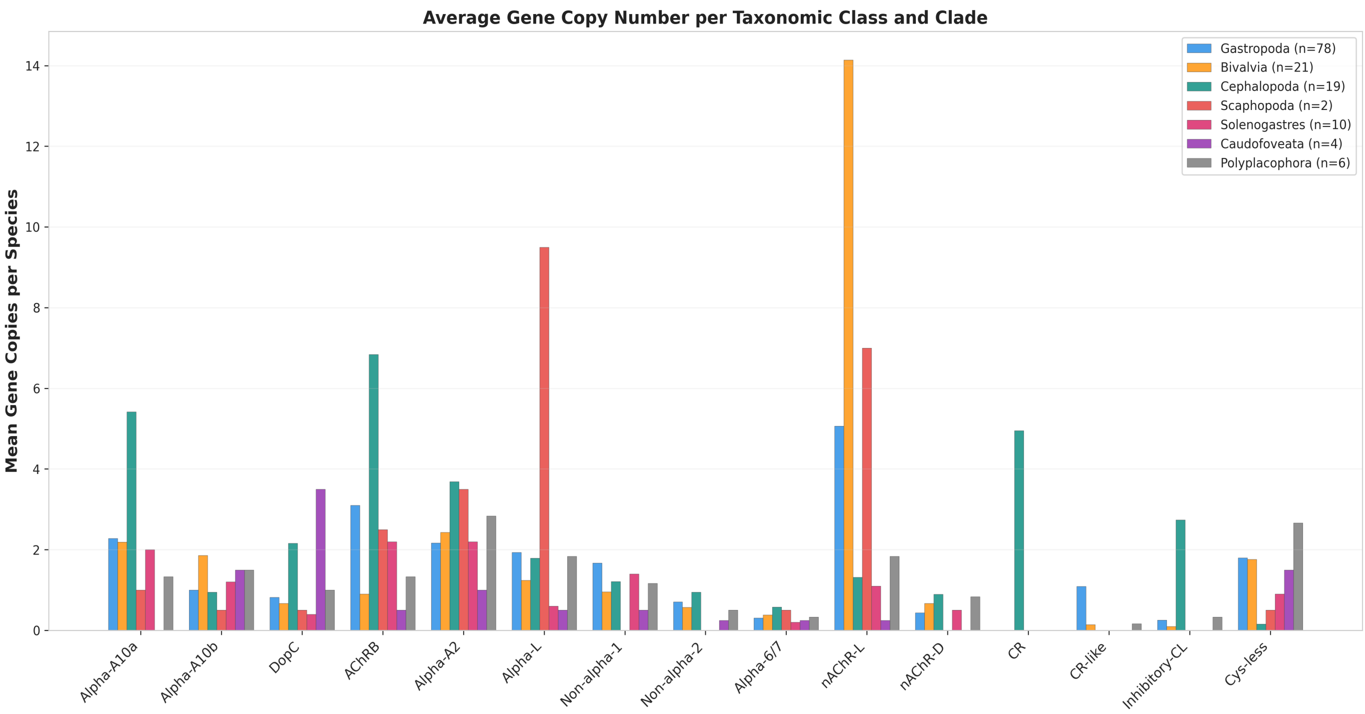


**Figure S3.** Average nAChR superfamily gene copy number per species across molluscan taxonomic classes and receptor clades. Bar heights represent the mean number of sequences assigned to each clade for species within each taxonomic class.

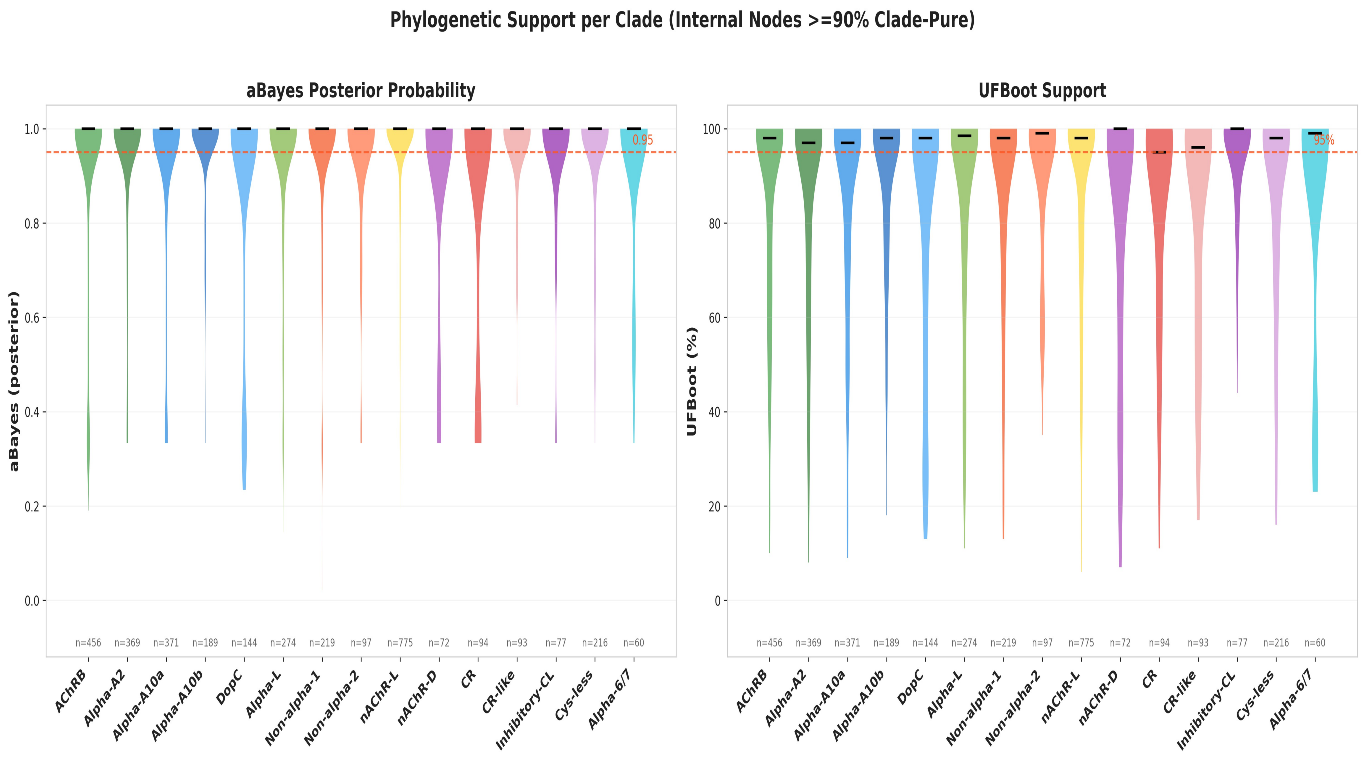


**Figure S4.** Phylogenetic support values for nAChR superfamily clades. (Left) Distribution of approximate Bayes posterior probabilities (aBayes) for internal nodes within each clade. (Right) Distribution of ultrafast bootstrap (UFBoot) support values. The analysis was performed using IQ-TREE2 with the Q.pfam+G8 substitution model and 1,000 ultrafast bootstrap replicates.

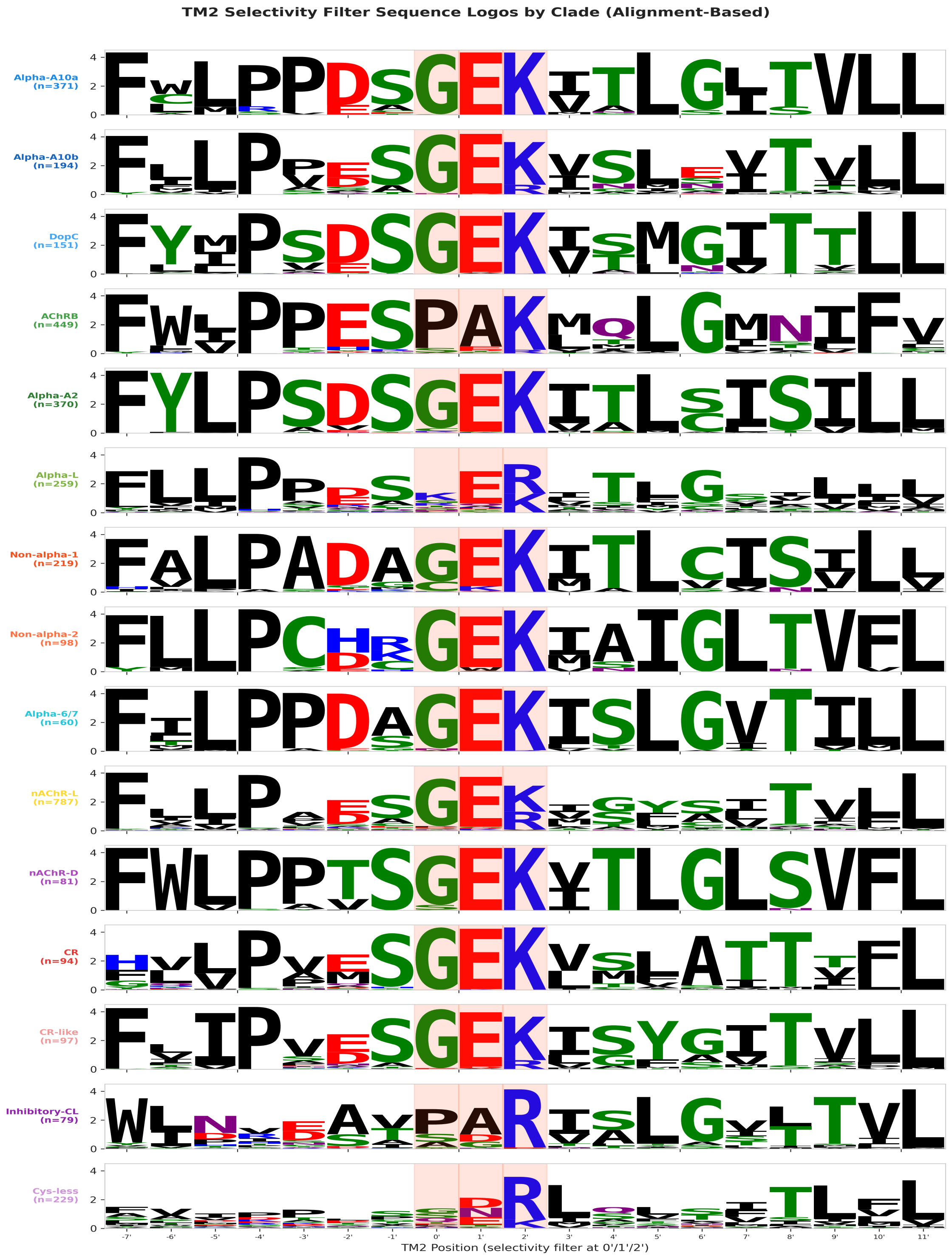


**Figure S5.** Sequence logos of the TM2 selectivity filter region across nAChR superfamily clades. Each logo represents the amino acid composition at the TM2 selectivity-determining positions for sequences assigned to each clade. The conserved GEK motif characteristic of cation-selective channels is prominent in most clades.

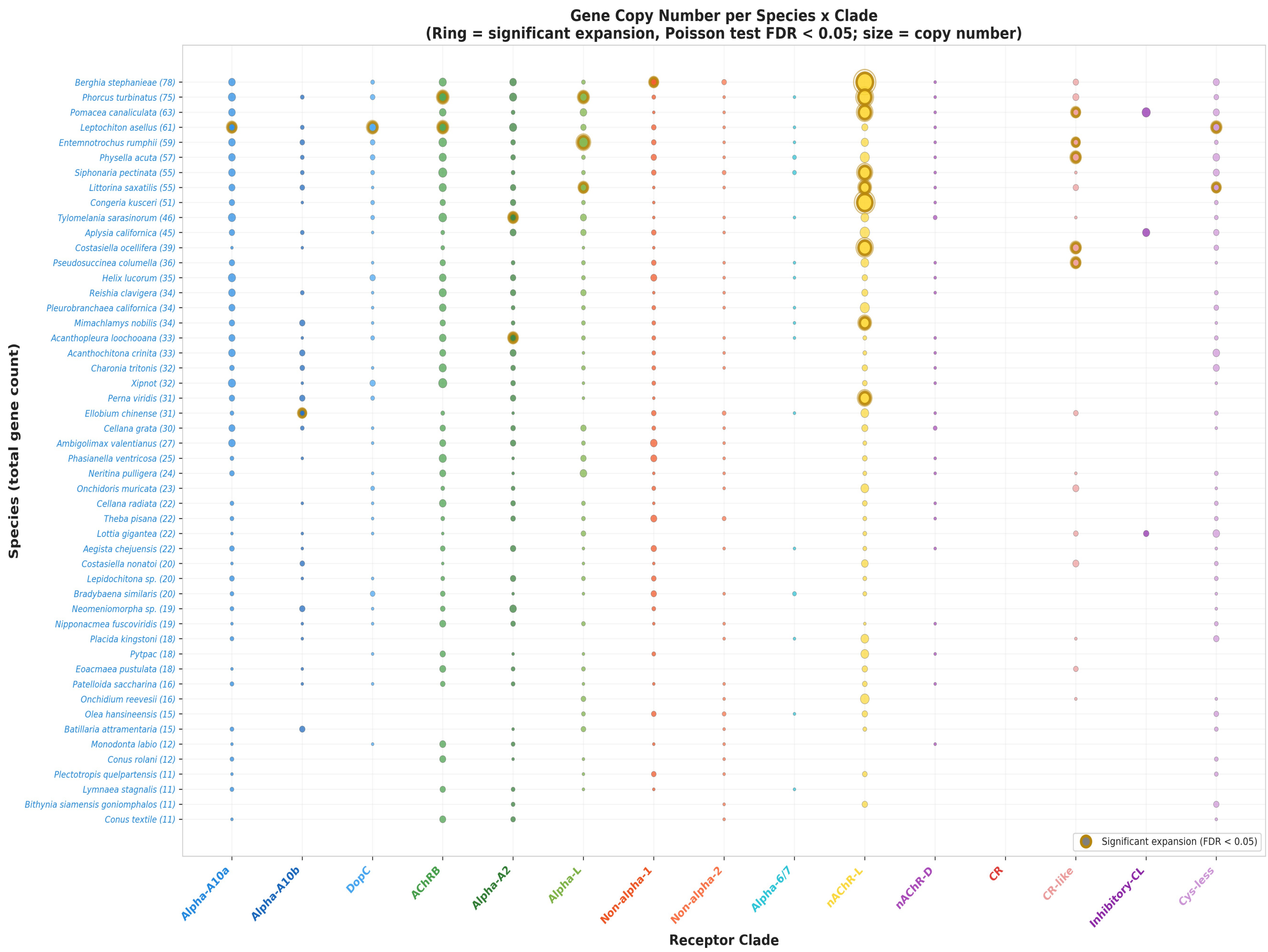


**Figure S6.** Gene copy number per species across nAChR superfamily clades. Each bubble represents the number of sequences assigned to a given clade for a given species, with bubble size proportional to copy number. Ringed bubbles indicate statistically significant expansions (Poisson test, FDR < 0.05).

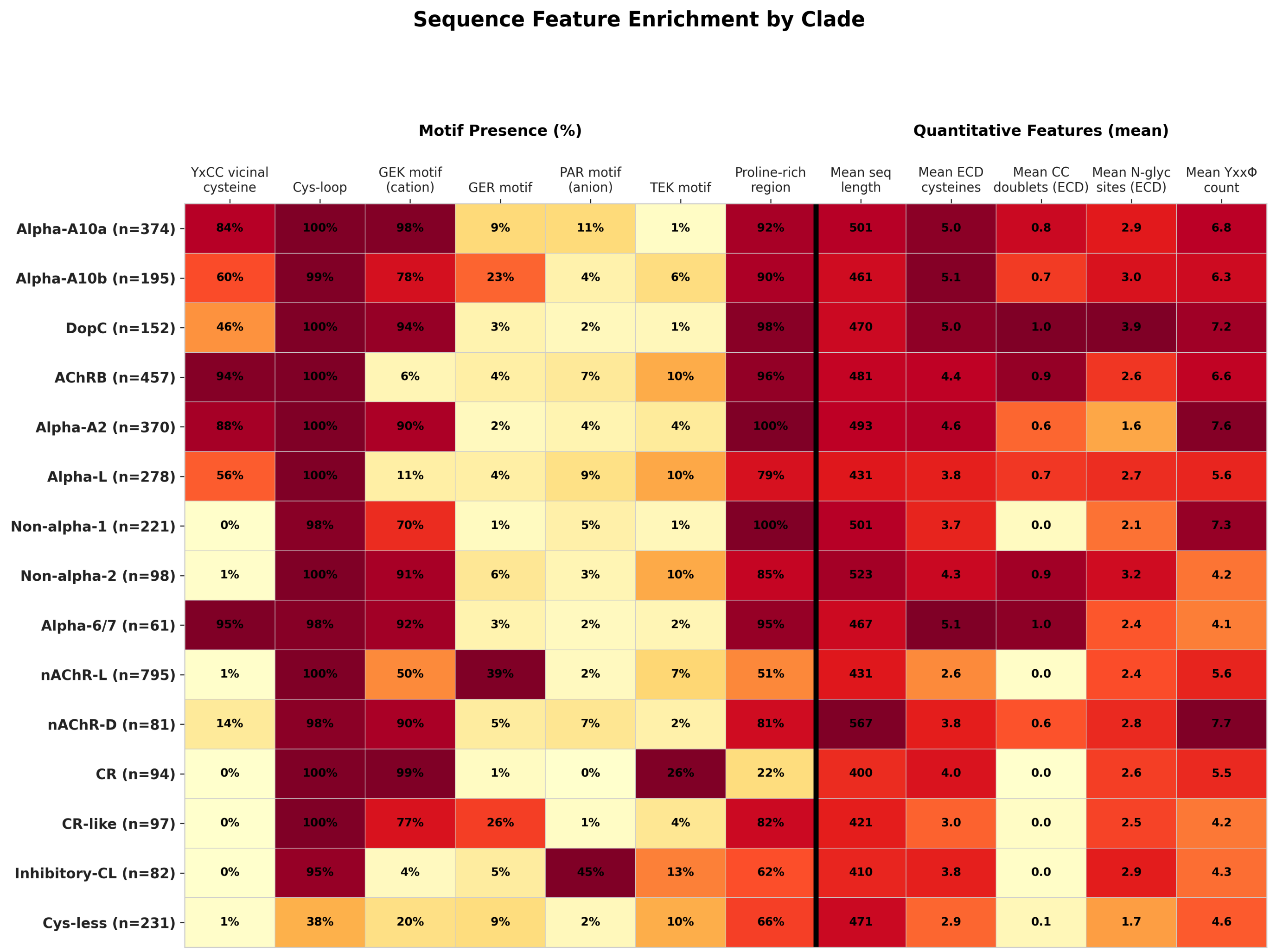


**Figure S7.** Sequence feature enrichment across nAChR superfamily clades. Heatmap showing the proportion of sequences within each clade possessing specific structural and functional motifs, including the YXCC vicinal cysteine (Cys-loop) motif, selectivity filter motifs, and quantitative features.

### Supplementary Methods

**Taxonomic Subsampling Strategy**

To ensure balanced taxonomic representation, a subsampling strategy was employed that retained all candidates from priority species (key model organisms including *Berghia stephanieae*, *Aplysia californica*, *Lymnaea stagnalis*, *Octopus bimaculoides*, and *Sepia officinalis*) while distributing remaining slots across taxonomic groups with maximum per-species and per-group caps to prevent overrepresentation of heavily sequenced lineages.

**Rogue Taxa Analysis**

A total of 125 sequences (3.5% of the dataset) were identified as rogue taxa and excluded from clade-level analyses. Rogues were detected based on long-branch attraction (63 sequences), assignment instability across bootstrap replicates (147 instances), monophyly-breaking placement (15 instances), and orphan block detection (181 instances). The majority of rogues originated from the nAChR-L (63/795, 7.9%) and Alpha-L (34/278, 12.2%) clades.

**Motif Extraction Pipeline**

Sequence features were extracted from aligned sequences using an automated pipeline that identified Cys-loop motifs (YXCC vicinal cysteine pairs), TM2 ion selectivity determinants (GEK, GER, PAR, and TEK motifs), signal peptide presence, N-glycosylation sites (N-X-S/T sequons), cysteine conservation patterns in the extracellular domain, and proline-rich regions.
